# Preparation of functional IgY that potently neutralizes HIV-1 in TZM-bl Cell line

**DOI:** 10.1101/2021.03.04.431083

**Authors:** Jinchen Wei, Yanqun Zhang, Yonglian Zeng, Yang Yang, Ronggan Liang, Chengqin Liang, Zhigang Dai, Songqing He, Shuwen Wu, Qing Xu

## Abstract

AIDS caused by HIV is one of the most serious public health challenges in the world. As we all know, Antiretroviral therapy (ART) is the most effective way to treat AIDS so far, however, forthe reasons of drug resistance, side effects, compliance, economy, limited its using widely. On the other hand, AIDS cannot be completely cured by ART. While the characterization of bnAbs (broadly neutralizing antibodies) in potent HIV neutralization provides considerable insight into HIV curing, it also can be used for passive immunotherapy or combination with ART for HIV-1 treatment. Here we report a novel technology to produce an neutralized activity bnAbs named HIV-1-IgY, which was extracted from the immunized Chicken egg by pNL4-3 virus antigens, and further purified using Water dilution and Salting out method. The specificity, titer and neutralizing activity of HIV-1-IgY was analyzed by Western blotting, ELISA and TZM-bl cell line evaluation system respectively. The results showed that theHIV-1-IgY has high neutralized activity to HIV in vitro; nearly 90% of HIV-1 were neutralized at 1.89μM in TZM-blsystem, which indicated that IgY may be a source of antibodies for AIDS prevention and treatment. Despite its needs to further consider and evaluate neutralized activity in-vivo and the potential mechanisms, Our data showed that we obtained an HIV-1-IgY which could effectively neutralize HIV-l IIIB virus in vitro.

## 1. Introduction

AIDS (Acquired immunodeficiency syndrome) is a public health problem worldwide, more than 2 million children and about 34.8 million adults had been infected (Frank et al., 2019). Antiretroviral therapy (ART) is standard scheme of AIDS treatment recommended by the FDA (Richman, 2001), however, it was restricted by drugs cost, potential long-term toxicities, and lifelong therapy (De Clercq, 2010; Kebede et al., 2017). Besides, ART is not an eradication strategy for HIV infection. Meanwhile broadly neutralizing antibodies (bnAbs) have attracted researchers’ attention forit powerful neutralized activity and benefits to guide vaccine design. Several antiviral activity experiments of broadly neutralizing antibodies had been performed in nonhuman primates. In course of prevention Simian Human Immunodeficiency Virus (SHIV) in non-human primates, individual administration of bnAbscouldinhibit SHIV effectively (Hessell et al., 2010; Hessell et al., 2009; Gautam et al., 2016). Therapeutic bnAbs have long been recognized as an effective and durable intervention strategy for AIDS suppressed.

Immunoglobulin Y (IgY) functions equivalent to immunoglobulin G (IgG) (Leslie and Clem, 1969). Despite functional homology among mammalian IgG and avian IgY, there are disparities in biochemical actions, molecular weight, and structure (Makoto et al., 1988). (1) IgY is remarkably stable in hypersaline environments and stabilizing reagents, which activity could remain 91% and 63% respectively, even under gastric conditions. Therefore, IgY can be administrated orally (Makoto et al., 1988; Rahman et al., 2013; Shin et al., 2013). (2) IgY neither activated the complement system nor interacted with rheumatoid factors in immune-assays(Carlander et al., 2000), it benefits for binding specificityand prevent non-specific inflammation(Dávalos-Pantoja et al., 2000; Larsson et al., 1993). (3) IgY can be produced in egg yolk and content abundant without sacrificing lives (Tini et al., 2002; Kovacs-Nolan et al,. 2005). For instance, one egg yolk contains approximately 100 mg IgY antibodies, which is much more than that isolated from immunized rabbits. For the other hands, the titer of IgY in egg yolk from immunized chickens had been demonstrated that significantly maintain high level for a long duration (Gassmann et al., 1990). Usage of IgY for treatment human and livestock disease have garnered researchers attention in past few decades, such as rotavirus-associated diarrhea, *Toxoplasma gondii* infection (Thu et al., 2017; Liou et al., 2010; Müller et al., 2015), There also reported finding an HIV-specific IgY, and test the neutralized activity through ELISA, indicated wide range uses of IgY. (Sudjarwo et al., 2014).

In this study, we reported how to prepare an functional anti-HIV-1 IgY (bnAbs) byusing HIV-infectious clone pNL4-3 as an HIV vaccine to immunize chincken, and further evaluated its neutralizing effect to HIV in HIV-l IIIB/TZM-bl system.

## 2. Materials and methods

### 2.1. Viral antigen preparation

The pNL4-3 plasmid which can release HIV-1 virus acquired from NIH in the United States and revived by the Virus Research Institute of Wuhan University.

1. Preparation of high purity pNL4-3 plasmid: The pNL4-3 plasmid was obtained by alkaline lysis and cesium chloride density gradient ultracentrifugation for 20 h. The super-helix plasmid ring was precipitated by syringe, then cesium chloride and ethidium bromide were pooled out by repeatedly extracting with chloroform. The plasmid was precipitated with isoamyl alcohol and dissolved in TE for storage.
2. pNL4-3 virus packaging: 293T cells (5×10^6^) were inoculated into 100 mm culture dish. After 24 h, pNL4-3 (10 μg) was used to transfect cells, collecting culture supernatants after 48 h and 72 h, storage at –80 °C.
3. Purification of pNL4-3 virus: The virus supernatant was mixed with 5×PEG8000 in a ratio of 4 to 1 at 4°C overnight and centrifugated at 4000 g for an hour at 4°C. Virus precipitate was washed thrice with phosphate-buffered saline (PBS), which was also used to resuspend the virus, quantify, sub-pack and store at −80 °C.

### 2.2. Immunization of laying hens

Specific pathogen-free (SPF) laying Leghorn hens, weight 1.5-1.7 Kg, 22 weeks of age, were provided by Hao Tai experimental animal breeding Co., Ltd. (Shandong province, China) and were fed at room temperature 20 to 25°C, humidity 50 to 60%, in sterile conditions. After a week of normal egg-laying, hens were numbered, divided into vaccination group and control group each for three hens. The HIV-pNL4-3 vaccine was mixed with an equal volume of Freund’s incomplete adjuvant and fully emulsified. The amount of antigen was quantified to 50-200 μg/mL and administrated 1mL/per hen through intramuscular injections on both sides of chicken wings and abdomen, once every seven days and repeated four times. The control group was given the same amount of saline and Freund’s incomplete adjuvant mixture in same way, collected eggs, numbered and preserved at 4°C every day.

### 2.3. IgY purification

Egg separator was used to separate yolk from whites, mixed egg yolk with pure water (equivalent to six yolk volumes), Stirring for 15 minutes then adjusting the pH to 5 (4.5-5.2). The mixture was kept at 4°C overnight and spun at 4000-10000 g for 40 min at 4°C, collecting the supernatant, 45% saturated ammonium sulfate was added to supernatant, mixed well and stand for 3hours at 4°C. The mixture was centrifuged at 4000-10000 g for 10 min at 4°C, Discard the supernatant and added equal volume of ddH_2_O. Then 13% sodium sulfate (mass-to-volume ratio) was added, mixed and kept still for 3hours. The mixture was centrifuged at 10000 g for 10 min at 4°C, the supernatant was discarded, and dissolved precipitate in 2 mL PBS and kept in dialysis membrane (50 KD cut off). The dialysis membrane was immersed in ddH_2_O with stirring, and exchanged ever per hour for 4-5 times, the last dialysis was carried out overnight. The final purified IgY liquid was stored at −20°C.

### 2.4. ELISA

The IgY titer of HIV-1 IgY was measured through indirect ELISA (Enzyme-Linked Immunosorbent Assay). HIV-1 gp41 was coated in plate wells ahead. Incubating the plate with serial dilutions of specific IgY at 37°C for an hour, PBS contain 0.05% Tween20 was used to wash the plate thrice. adding rabbit anti-chicken IgY which HRP-conjugated secondary antibodies (Promega, Madison, WI, USA) in the dilution of 1:5000 and incubated at 37°C for an hour. TMB color reaction was used to evaluate the tilter of HIV-1 IgY after washed thrice. Finally, The absorbance was measured at 450 nM.

### 2.5. SDS-PAGE and immunobloting assay

The purity and specificity of IgY were measured by immunobloting assay. Briefly, the purified IgY were run on 10% SDS-PAGE and subsequently stained by Coomassie brilliant blue to analyze the purity. For the specificity of IgY assay, pNL4-3 virus lysate was loaded as antigens and separated on 10% SDS-PAGE, then transferred to nitrocellulose, blocked with 5% skim milk, and incubated with IgY at 4°C overnight. TBST washed three times ever 5 min, incubated with anti-IgY secondary antibody (Promega Madison, USA) for an hour, detected the bands through ECL-based method.

### 2.6. Neutralization assay

1.5 × 10^3^ TZM-bl cells were seeded into48 well plate and cultured overnight. Then TZM-bl cells were infected with an HIV-l IIIB virus at 0.1 mol. At the same time, IgY was diluted into a concentration gradient and added to corresponding wells, cultured 2 h at 37°C. Then replaced the culture medium with fresh medium without virus and IgY. After continuous cultured at 37°C for 48 h, the culture medium was discarded, added 30μL cell lysis buffer into each well and incubated at room temperature for 15-20 min. The samples were then treated with Promega luciferase substrate kit according to the instructions, and the activity of the reporter gene was detected (GloMaxr 20/20 luminometer, Promegacompany). The percentage of IgY inhibitory activity was calculated according to the following formula: [1-(E-N)/(P-N)] X 100%, wherein negative control: N, positive control: P and the test group: E. The effective concentration for 50% inhibition (EC_50_) of IgY was calculated by CalcuSyn software.

### 2.7 Statistical analyses

For neutralizing activity of IgY was compared using the unpaired t test in Prism (GraphPad). Statistical significance was assessed at p<0.05 for all comparisons.

## 3. Results

### 3.1. The purity and quantify of HIV-1 IgY

To measure the purity of IgY, SDS-PAGE was carried out, as showed in Fig 1, two major bands (68 KD and 30 KD) and two minor bands (50 KD and 42 KD) are observed, indicatingthat HIV-1 IgY contained 68 KD HC (heavy chain) and 30 KD (light chain) (Fig. 1). For analyzing the HIV-1 IgY antibody by gel imaging system software, the purity of HIV-1 IgY was up to 85%.

**Fig.1.**
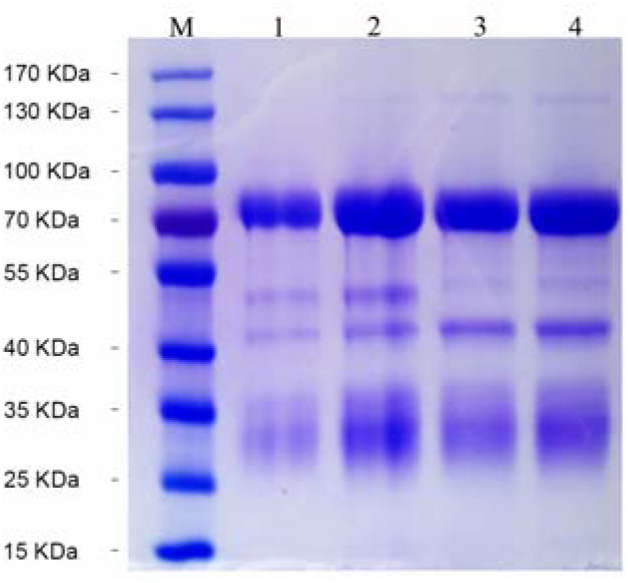
The purity of HIV-1 IgY. M, Markers; Lane 1 and Lane 2 were the same sample with different load volume (10μL with Lane 1, 20μL with Lane 2 respectively), IgY post-immunization with pNL4-3 virus antigen; Lane 3 and Lane 4: IgY post-immunization with Freund’s incomplete adjuvant. Lane 3 and Lane 4 were also the same sample with different load volume (10μL with Lane 3, 20μL with Lane 4 respectively).

Pre-immunized eggs contain 0.74 ± 0.056μM IgY in yolk. After immunized with PNL4-3, the concentration of IgY markedly increased (8.61 ± 0.44 μM) from second week and reached a maximum of 14.5μM in week11th. Then, a gradual decrease was observed, in week 13th, the content of IgY decreased to 5 ± 0.22 μM (Fig. 2).

**Fig.2.**
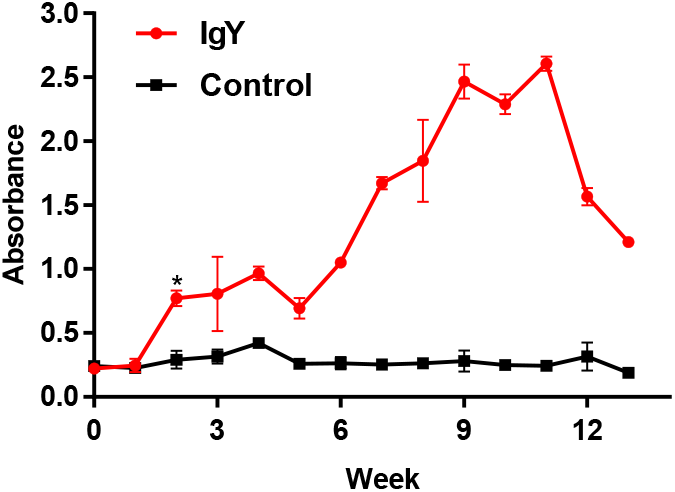
The titer of IgY after Immunization. After injected with pNL4-3 virus antigen in chicken at 0 week, the chicken immune system response quickly from second week. After 4 times immunization, there was a noticeable increase at 5th week and reached maximum at 11th week, then began to decrease.

### 3.2. The Specificity and neutralization activity of HIV-1 IgY

HIV-1 (from PNL4-3 plasmid) was subjected to SDS-PAGE, transferred to a nitrocellulose filter and probed with HIV-1-IgY for both immunized and non-immunized IgY. The HIV-1 IgY antibody was found to interact with 41KD and 120KD bands of HIV-1 (Fig. 3), and the neutralization activity of HIV-1IgY was analyzed through neutralization assay in vitro. As expected, HIV-1 IgY supplementation inhibited HIV-l IIIB virus proliferation dose-dependently, which inhibited 90% HIV-l IIIB virus at 1.89μM, similarly to AZT Zidovudine, indicating a broad neutralization activity of HIV-1 IgY (Fig. 4).

**Fig.3.**
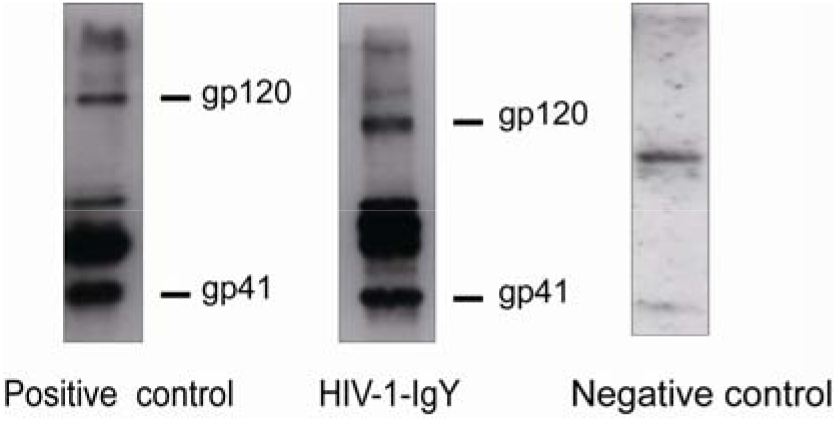
The ability of HIV-1 IgY to interact with HIV. The HIV antigens were load and separated by SDS-PAGE. And HIV-1-IgY was used as primary antibodies. Results showed HIV-1-IgY could specifically recognize the envelope protein gp41 and gp120 of HIV-1, while the negative control group had no immune reactivity with antigens.

**Fig.4.**
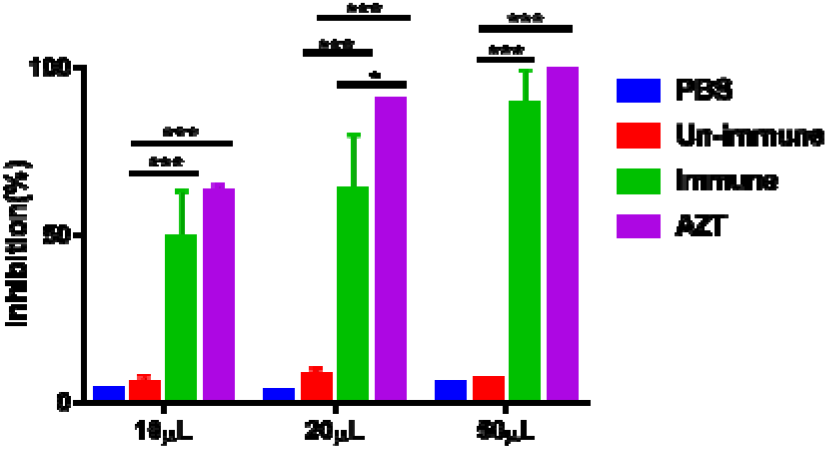
Neutralizing activity comparison of HIV-1-IgY and AZTin TZM-bl Cell line. The results showed IgY is HIV-1 specific, could restrict the expression of luciferase in TZM-bl cells which pNL4-3 virusinfected. For the 10, 20 and 50μL HIV-1 immune groups, the IgY concentrations were 0.38μM, 0.76μM, 1.89μM, respectively. The corresponding inhibition rates (50%, 64%, and 90%) showed a dose-response relationship. The corresponding inhibition rates of IgY in the pre-immune control group were 10%, 12%, and 12%, respectively. 6 for all groups. The difference between immune groups and un-immune group was statistically significant (*P* < *0.001*).

## 4. Discussion

Although HAART is still an effective therapy for AIDS, developing potent bnAbs that elicits reliable, broad neutralization is also important. Different from small molecular drugs, bnAbs can neutralize the HIV and kill the infected cells. However, to elicitreliable HIV bnAbs had not been tried by immunogen because of variable antigenic diversity of Env and glycan shield for HIV (Walker et al., 2011; Doria-Rose et al., 2014). Therefore, few antibodies respond to HIV in mammal, except cow and llama. As previously described, repeated immunization with non-well-ordered Env trimer in cow and llama could generate potent and broad neutralization HIV-1 antibodies (Sok et al., 2017; McCoy et al., 2012). However, few other studies also showed that birds can produce bnAbs, although their biological effectiveness remains unknown.

In this study, we characterized rapid and long-term methods to produce bnAbs in chicken eggs, which could potently neutralize HIV-1 in TZM-bl cell line. By multiple intramuscular administrations, we detected high level of bnAbs in egg yolk after two weeks (OD>1.009 ± 0.05), faster than other methods which described in previous study in cows (after 4 weeks) and llama (after 2 months) (Sok et al., 2017; McCoy et al., 2012); bnAbs remained high, peaked at 9^th^ week and sustained until 11^th^ week. Subsequently, we analyzed the interaction between bnAbs and Env by immunobloting assay, HIV-1 IgY could bind to virus Env, suggesting the specificity to HIV of HIV-1IgY. Then we explored the biological activity of HIV-1IgY in vitro and almost 90% TZM-bl/HIV-1 were inhibited by HIV-1-IgY 1.89μM, similar to Zidovudine. Besides, we used dialysis followed by ammonium sulfate precipitation to purityHIV-1 IgY, which is simple and efficient for mass production, considering the safety applications and sustainable production (Mulvey et al., 2011; Vega et al, 2012). HIV-1-IgY (bnAbs) antibodies could either bind or potently neutralize HIV-1 in TZM-bl cell line. Other advantages such as stable of IgY provide a potential basis for HIV-1IgYdevelopingto a clinical agent in the future. For the other hand, HIV-1-IgY (bnAbs) may be combined with HARRT in HIV-1 treatment, or serve as prophylactic and diagnostic agent for HIV infection.

## Acknowledgments

This work was supported by the National Natural Science Foundation of China (No. 81560574 and 31560100). The work was also funded by the Molecular Medicine of Liver Injury and Repair Collaborative Innovation Center and Guangxi Bagui Scholars project. We thank Professor Zhi-Kai Dai, researcher Nin Tan, Li-Ya Li (Guilin Medical University), Qiu-Juan Chen and Professor Xing-Can Shen (Guangxi Normal University) for the meaningful exploratory work in the early stage of the study.

